# Targeted induction of *de novo* Fatty acid synthesis enhances MDV replication in a COX-2/PGE_2α_ dependent mechanism through EP2 and EP4 receptors engagement

**DOI:** 10.1101/323840

**Authors:** Nitish Boodhoo, Nitin Kamble, Benedikt B. Kaufer, Shahriar Behboudi

## Abstract

Many viruses alter *de novo* Fatty Acid (FA) synthesis pathway, which can increase availability of energy for replication and provide specific cellular substrates for particle assembly. Marek’s disease virus (MDV) is a herpesvirus that causes deadly lymphoma and has been linked to alterations of lipid metabolism in MDV-infected chickens. However, the role of lipid metabolism in MDV replication is largely unknown. We demonstrate here that infection of primary chicken embryonic fibroblast with MDV activates *de novo* lipogenesis, which is required for virus replication. In contrast, activation of Fatty Acid Oxidation (FAO) reduced MDV titer, while inhibition of FAO moderately increased virus replication. Thus optimized virus replication occurs if synthetized fatty acids are not used for generation of energy in the infected cells, and they are likely converted to lipid compounds, which are important for virus replication. We showed that infection with MDV activates COX-2/PGE_2α_ pathway and increases the biosynthesis of PGE_2α_, a lipid mediator generated from arachidonic acid. Inhibition of COX-2 or PGE_2α_ receptors, namely EP2 and EP4 receptors, reduced MDV titer, indicating that COX-2/PGE_2α_ pathway are involved in virus replication. Our data show that the FA synthesis pathway inhibitors reduce COX-2 expression level and PGE_2α_ synthesis in MDV infected cells, arguing that there is a direct link between virus-induced fatty acid synthesis and activation of COX-2/PGE_2α_ pathway. This notion was confirmed by the results showing that PGE_2α_ can restore MDV replication in the presence of the FA synthesis pathway inhibitors. Taken together, our data demonstrate that MDV uses FA synthesis pathway to enhance PGE_2α_ synthesis and promote MDV replication through EP2 and EP4 receptors engagement.

## Author summary

Alteration of lipogenesis can be beneficial for viruses by providing energy for the host cells, and creating the required microenvironment for virus entry, assembly and egress. Disturbances of lipid metabolism in Marek’s disease virus (MDV) infected chickens are thought to contribute to viral pathogenesis. However, the role of the lipid metabolism in MDV replication remains unknown. Here, we demonstrate that infection of primary chicken cells with MDV activates fatty acid synthesis pathway and increases lipogenesis. Interestingly, optimized virus replication occurs when the synthetized fatty acids are not used for generation of energy, but instead utilized to promote the biosynthesis of PGE_2α_. We demonstrate that PGE_2α_ increases MDV replication through engagements of EP2 and EP4 receptors, which are upregulated in MDV infected cells. Taken together, we concluded that MDV-induced FA synthesis pathway leads to an increased PGE_2α_ synthesis which enhances MDV infectivity through EP2 and EP4 receptors on MDV-infected cells.

## Introduction

Despite their great diversity, viruses are highly dependent on host cell factors to facilitate maximal viral replication. Elucidating the mechanisms underlying hijacking of host cell metabolism by viruses can identify key determinant factors for virus replication which consequently contribute to an altered physiological homeostasis and disease susceptibility. All herpesviruses including Marek’s Disease Virus (MDV) encode metabolic enzymes in their genomes which are involved in their pathogenesis [1–4]. MDV is a highly oncogenic herpesvirus and the etiologic agent for Marek’s disease (MD). Upon infection of the host, the virus initially replicates in B and T cells. Subsequently, the virus establishes latency predominantly in CD4^+^ T cells, allowing the virus to persist in the host for life [5]. [5]. In addition, latently infected CD4^+^ T cell can undergo neoplastic transformation resulting in deadly lymphomas[5].

A diverse range of human enveloped viruses including human cytomegalovirus (HCMV) [6], herpes simplex virus 1 (HSV-1) [7], hepatitis C virus (HCV) [8], dengue virus (DENV) [9], Kaposi’s sarcoma-associated herpesvirus (KSHV), human immunodeficiency virus (HIV) [10], vaccinia virus (VACV) [11] and West Nile virus (WNV) [12], have been shown to selectively modulate FA synthesis pathway to support virus replication. In this process, acetyl-CoA is converted to malonyl-CoA and subsequently to palmitate. The first step towards FA synthesis is the conversion of citric acid into acetyl-CoA by direct phosphorylation of ATP-citrate lyase (ACLY). The subsequent committed step involves the conversion of acetyl-CoA into malonyl-CoA by acetyl-coA carboxylase (ACC), a process modulated by HCMV [13] and HCV [14]. ACC is the rate-limiting step and contributes to cholesterol synthesis. The final step involves the committed elongation by utilizing both acetyl-CoA and malonyl-CoA coupled to the multifunctional fatty acid synthase (FASN) to make palmitic acid. DENV [9], WNV [12], and HCV [15] have been shown to preferentially enhance FASN activity and palmitic acid synthesis.

Palmitic acid contributes to several key biological functions such as fatty acid oxidation (FAO), β-oxidation in mitochondria, post-translational modification (palmitoylation) of proteins or elongation of fatty acid chains to generate a diverse repertoire of very long chain fatty acids (VLCFA). This diverse role for the utilization of palmitic acid has been demonstrated in HCV [8], VACV [11], modified vaccinia Ankara (MVA) [16], HCMV [17,18], KSHV [19, 20], respiratory syncytial Virus (RSV) [21], and Epstein-Barr Virus (EBV) [22]. Both HCV [8] and vaccinia virus [11] are highly dependent on mitochondrial β-oxidation to support infection. On the other hand, recent data suggest that VLCFA are essential for HCMV replication [23]. VLCFA contribute to biosynthesis of Arachidonic acid (AA), which can be further reduced by cyclooxygenase-1 (COX-1) and inducible COX-2 to make prostaglandin E_2α_ (PGE_2α_), a potent eicosanoid and immune modulator, modulating the functions of natural killer (NK) cells [24], macrophages [25], and T cells [26]. Induction of PGE_2α_ biosynthesis has been observed following infections with HCMV [17, 18], KSHV [19, 20], RSV [21], Influenza A virus (IAV) [27] and MVA [16]. Moreover, a direct association between induction of COX-2 activity and enhancement of HCMV and KSHV replication has been reported [17, 20]. For example, PGE_2α_ inhibits macrophages recruitment to the lungs, reduces type | IFN production and modulated antiviral immunity in IAV infection. Intriguingly, inhibition of COX-2/PGE_2α_ pathway protected mice against IAV [27] and rescued the inhibitory effects of soluble factors released by MDV-transformed T cells on T cell proliferation [28]. Collectively, these studies suggest that induction of FA synthesis has major implications on virus replication and immune evasion and dissemination in the host.

Previously, atherosclerotic plaque formation has been reported in chickens infected with pathogenic MDV [29]. Vaccination with herpesvirus of turkey (HVT; MDV serotype 3) prevented the development of atherosclerotic plaques [30]. Lipid analysis of the arterial smooth muscles (ASM) from MDV infected birds revealed a significant increase in non-esterified fatty acids (NEFA), cholesterol, cholesterol esters, squalene, phospholipids and triacylglycerol. Furthermore, excess lipids biosynthesis triggered cellular deposition in organelles termed lipid droplets [29, 30]. Despite these intriguing observation, it remained elusive if the fatty acid metabolism is altered in MDV infected cells and how MDV causes these alterations in the host.

In the present study, we investigated if FA synthesis and FA derivatives are induced in MDV infected cells. We could demonstrate that MDV infection induce *de novo* FA synthesis and upregulation of genes involved in FA synthesis and FAO. Using small pharmacological inhibitors and/or addition of exogenous fatty acids, we could show that FA synthesis pathway is essential to support MDV infection in cultured cells. Short chain FAs contributed to LCFA synthesis which can be stored in lipid droplets. However, the virus replication was not dependent on the ATP production from utilization of short chain FAs by FAO. MDV infection induced COX-2 expression and PGE_2α_ synthesis, and this process was dependent on activation of FA synthesis pathway by MDV. Taken together, our results demonstrate that the major role of FA synthesis to support MDV replication is to activate PGE_2α_/EP2 and PGE_2α_/EP4 signalling pathways.

## Results

### MDV infection increases lipogenesis

To determine if MDV infection affects the lipid metabolism, we performed a lipidomic analysis on mock and MDV infected primary chicken embryo fibroblasts (CEFs) at 48 and 72 hours post infection (hpi). The analysis revealed that 15 lipid metabolites were increased and 9 metabolites were decreased in infected cells at 72 hpi (Fig 1A). To provide a better visualization of the metabolic changes, we mapped the altered fatty acids (FA) and their derivatives onto FA synthesis pathway at 48 (Fig 1B i) and 72 hpi (Fig 1B ii). At 72 hpi, increased levels of palmitic acid (p = 0.01), stearic acid (p = 0.03), Oleic acid (p = 0.01), Nervonic acid (p = 0.04), Mead acid (p = 0.0028) and Docosahexanoic acid (p = 0.001) were observed. In contrast, Arachidic acid and Lignoceric acid were significantly reduced in MDV infected cells (p = 0.001). Alternatively, palmitic acid can be utilized for phospholipid biosynthesis. Our data demonstrate that phosphatidylethanolamine (PE) synthesis is also upregulated in the infected cells at both 48 (p = 0.0005) and 72 (p = 0.001) hpi. Similarly, arachidonic acid (AA) was also significantly increased at both 48 (p = 0.0031) and 72 hpi (p = 0.0028). Taken together, our data demonstrate that MDV infection alters lipid metabolism and increases synthesis of palmitic acid and long chain fatty acids.

**Fig 1:**
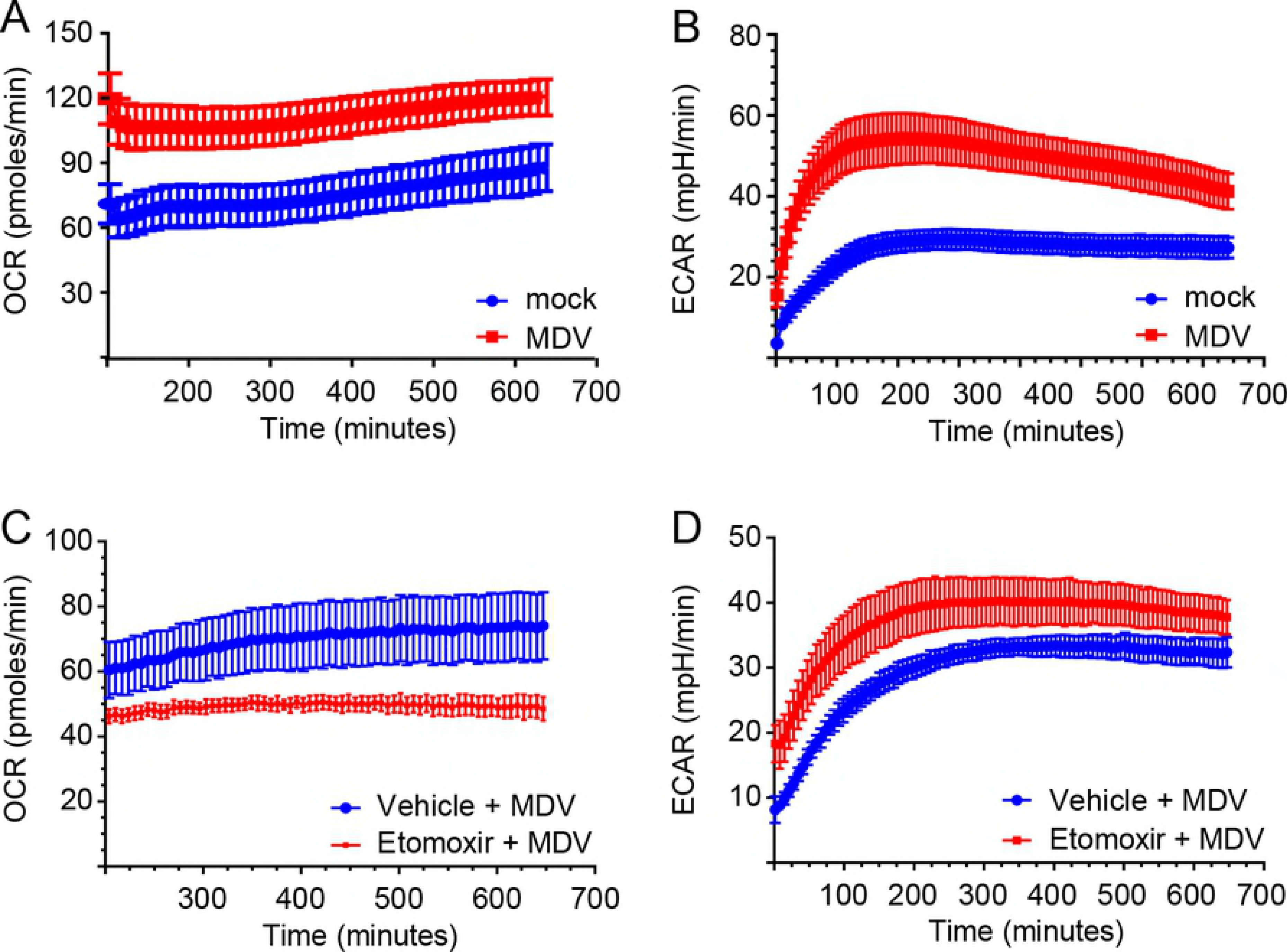
Induction of *de novo* Fatty acids synthesis in MDV infected CEFs. Metabolomics analysis of relative levels of lipid metabolites from mock-(control) and MDV-infected (RB1B) CEFs are shown at 48 hpi and 72 hpi. **(A)** Box and whisker plots showing minimum and maximum relative levels of named fatty acids and long chain fatty acids LCFAs) either significantly altered or where no changes were observed as a result of MDV infection. **(B)** Schematic summary of major metabolites outlining preferential utilization of fatty acids for LCFA and phospholipid synthesis by comparing (i) 48h and (ii) 72h post MDV infection. Arrows indicate 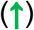 increase, 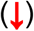 decrease or (→) no change in the respective metabolites to demonstrate flux through pathways. Non-parametric Wilcoxon tests (Mann-Whitney) was used to assess normal distribution and test significance with the results shown as mean ± SD **(A)**. * (p = 0.01) and ** (p = 0.001) and *** (p = 0.0005) indicates a statistically significant difference compared to control. NS indicates no significant difference. The experiment was performed in biological triplicates with six technical replicates per biological replicates

### MDV replication requires Fatty acid synthesis

To examine the role of *de novo* lipogenesis in the increased levels of fatty acids in MDV infected cells, we performed gene expression analyses of the cellular enzymes involved in FA synthesis pathway using RT-PCR (Fig 2A). Both acetyl-CoA carboxylase (ACC) and Fatty acid synthase (FASN) were highly upregulated at 72 hpi (Fig 2B). To determine the role of fatty FA synthesis pathway in MDV replication, we used selective inhibitors of ACLY (SB 204990), ACC (TOFA) and FASN (C75) (Fig 2A). Non-toxic concentrations of the inhibitors were determined based on viability (S1 Fig) and confluency of the treated CEFs. CEFs were infected with MDV in the presence of the pharmacological inhibitors with/out the downstream metabolites (malonyl-CoA and palmitic acid) and MDV titers were quantified by plaque assays at 72 hpi. The ACLY inhibitor (SB 204990) did not alter MDV replication (Fig 2C), suggesting that the conversion of citrate to Acetyl-CoA is not important for the virus replication. While, inhibition of ACC (TOFA; 1.54 μM) and FASN (C75; 5.9 mM) significantly impaired MDV replication by 27 (Fig 2D) and 28 folds (Fig 2E), respectively. Treatment of MDV-infected CEFs with a combination of TOFA (0.77 μM) and C75 (4.5 μM) decreased (p = 0.0043) the transcripts of ACC (Fig 2F) and FASN (Fig 2G) as determined using RT-PCR. Nest we examined if the downstream metabolites can rescue virus replication in the presence of the inhibitors. As shown above, SB 204990 did not alter virus titer even when malonyl-CoA was added to the culture (Fig 2H).

**Fig 2:**
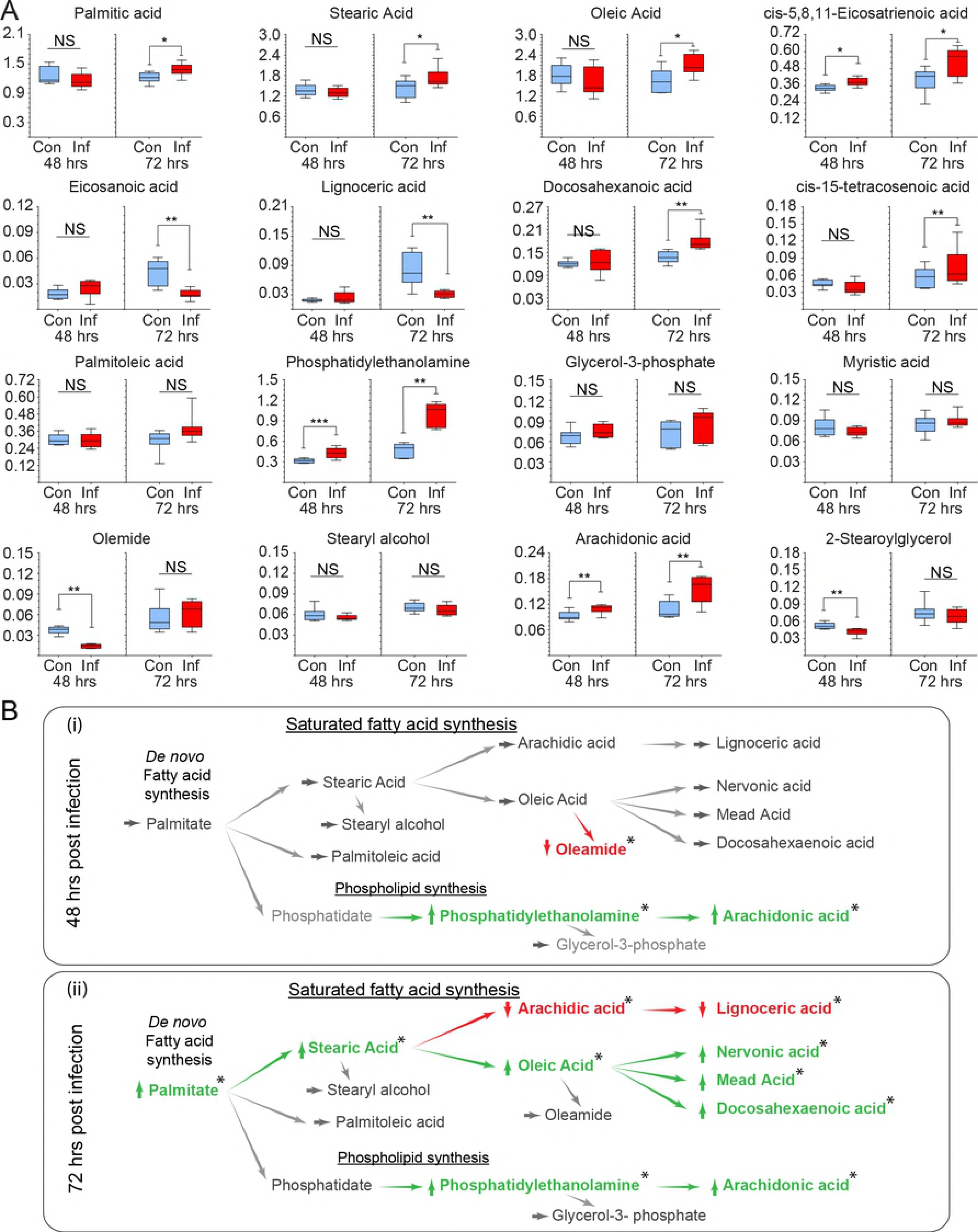
Fatty acid synthesis is required for MDV infection. A pathway interference approach to dissect the role of lipids in MDV replication. **(A)** Schematic FA synthesis pathway highlighting the relevant pharmacological inhibitors (red) and the respective enzymes (yellow box) as well as metabolites (green box) studied within FA synthesis pathway. **(B)** Fold change gene expression in CEFs mock-infected or MDV-infected are shown at 24, 48 and 72 hpi. Analysis of MDV viral titer (PFU/ml) in infected CEFs in the presence of **(C)** SB 204990 (1.28, 2.56 and 3.85 μM), **(D)** TOFA (0.03, 0.077, 0.154, 0.31, 0.77 and 1.54 μM) and **(E)** C75 (0.393, 1.97, 3.93 and 5.9 μM). Box and whisker plots demonstrating relative fold change in mRNA of **(F)** ACC and **(G)** FASN in MDV-infected CEFs cells treated with vehicle (Cont.) or TOFA+C75 (T/C) at 72 hpi. Analysis of MDV viral titer (PFU/ml) in infected CEFs in the presence of **(H)** malonyl-CoA with SB-204990, **(I)** malonyl-CoA with TOFA or **(J)** Palmitic acid (5, 12.5, 20 and 25 μM) **(K)** Palmitic acid with C75 (5.9 μM). **(L)** MDV (RB1B) genome copy number per 10^4^ cells (MEQ gene with reference ovotransferrin gene) in the presence of SB 204990 (3.85 μM), TOFA (1.54 μM) or C75 (5.9 μM). *** (p = 0.0002) **** (p < 0.0001) indicates a statistically significant difference compared to control. NS indicates no significant difference. $ symbol indicates vehicle treated cells. All viral titer experiments were performed in 6 replicates and data is representative of 3 independent experiments.

Interestingly, addition of malonyl-CoA restored MDV titers, which was inhibited by TOFA (Fig 2I). Similarly, palmitic acid restored virus replication in the presence of C75 (Fig 2K), while it did not alter MDV titers on its own (Fig J). We also demonstrated that TOFA and C75, but not SB 204990, reduced gene copy numbers as determined using qPCR (Fig 2L). Taken together, the result demonstrate that blocking ACC and FASN decreases MDV replication and reveal that palmitic acid is a key metabolite required for MDV infection.

### Fatty acid are utilized to generate energy in the MDV-infected cells

To determine if fatty acids oxidation increases ATP synthesis, we measured oxygen consumption rate (OCR) and extracellular acidification rate (ECAR) in MDV-infected cells over time using a Seahorse Bioscience XFp analyzer (Seahorse Bioscience). Thereby, OCR and ECAR correspond to β-oxidation of fatty acids and glycolysis for ATP production, respectively. Linear regression analysis of the OCR and ECAR revealed a significant increase in MDV-infected compared to mock-infected cells (Fig 3A and 3B), suggesting that virus infection increases ATP production via both β-oxidation of fatty acid and glycolysis. To assess if MDV replication relies on import and utilization of palmitic acid within mitochondria to generate ATP, CEF cells were treated with a vehicle or etomoxir, the pharmacological inhibitor of carnitine palmitoyltransferase | (CPT1a). Etomoxir did not modulate β-oxidation of fatty acids in the non-infected cells (S2 Fig). In contrast, etomoxir significantly reduced the OCR in MDV-infected cells (Fig 3C). Intriguingly, treatment of MDV-infected cells with etomoxir increased the ECAR rate at all time points (Fig 3D), indicating that glycolysis can replace energy source provided by FAO in the MDV-infected cells.

**Fig 3:**
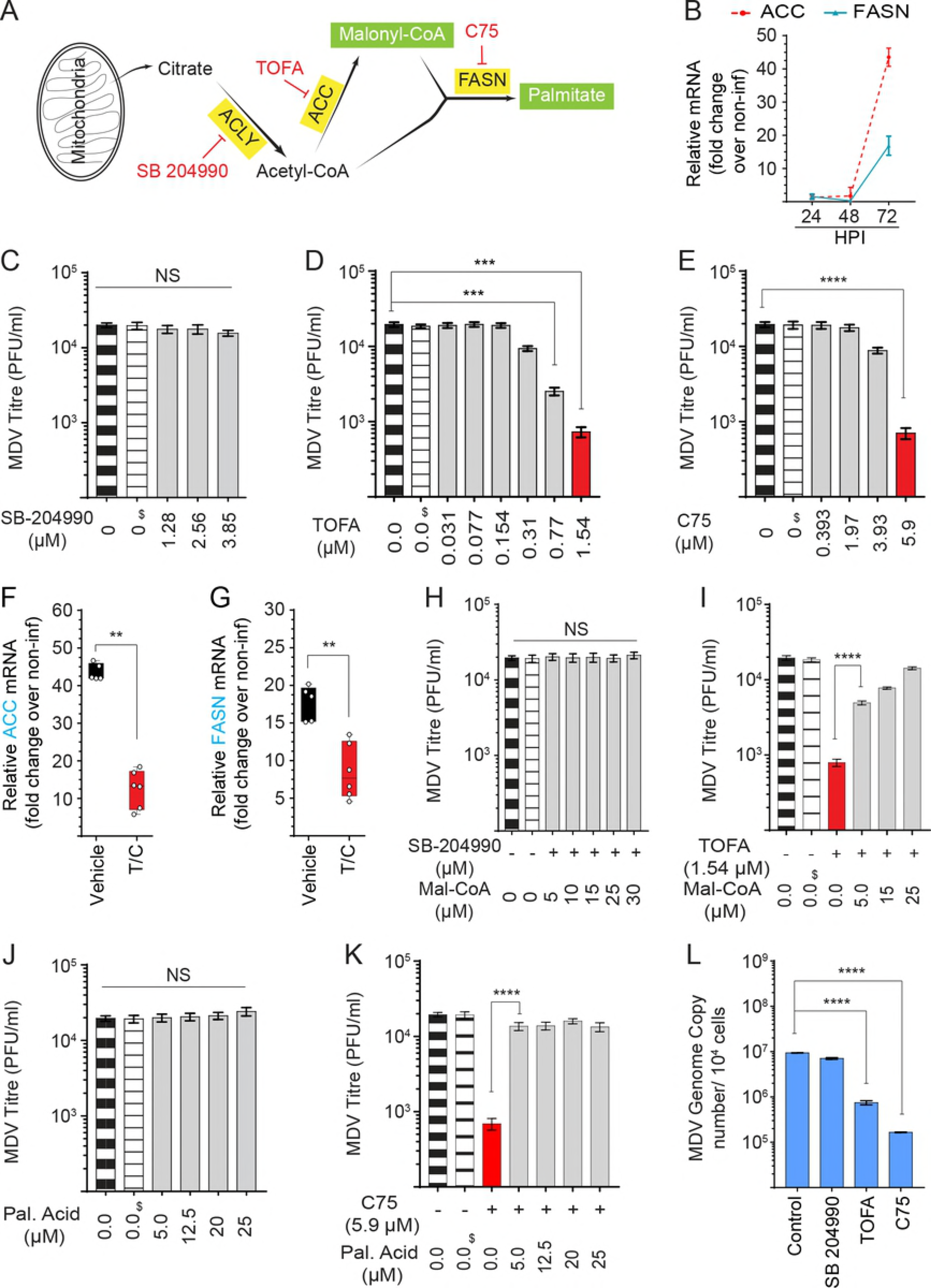
MDV infected cells can also utilize non-mitochondrial sources of ATP. **(A)** Oxygen consumption rate (OCR; pmoles/min) and **(B)** Extracellular acidification rate (ECAR; mpH/min) are shown for the mock-infected and MDV-infected CEFs. MDV-infected CEFs were treated with etomoxir (4.42 μM) or vehicle during the first 12 h post mock infection, and **(C)** OCR **(D)** ECAR were determined using the Seahorse XFp. All experiments were performed in triplicates and data is representative of 3 independent experiments.

### Higher virus replication occurs if the synthetized fatty acids are not used for generation of energy

To investigate if the ATP generated by β-oxidation are required for MDV replication, we inhibited CPT1a responsible for fatty acids transport from cytoplasm into mitochondria for β-oxidation using non-toxic concentrations of the physiological inhibitor of CPT1a, malonyl-CoA, and the pharmacological inhibitor, etomoxir. To activate β-oxidation, we used nontoxic concentrations of Clofibrate, a PPARα ligand which activates β-oxidation, in this study (Fig 4A). Gene expression analysis revealed a moderate up-regulation of CPT1a, acyl-CoA dehydrogenase long chain (ACADL) and lipoprotein lipase (LPL) in the MDV-infected CEFs using RT-PCR at 72 hpi (Fig 4B), indicating that FAO is activated in MDV infected cells. To examine the role of β-oxidation in MDV-replication, CEFs were infected with MDV in the presence of exogenous malonyl-CoA, etomoxir or Clofibrate, and viral titer was quantified at 72 hpi. Clofibrate significantly reduced MDV replication (Fig 4C). In contrast, neither malonyl-CoA (Fig 4D) nor etomoxir (Fig. 4E) had no inhibitory effects on virus titers, suggesting that the ATP generated by β-oxidation is not required for MDV replication. Interestingly, even a minor but significant increase in viral yield was observed in the cells treated with etomoxir or malonyl-CoA. Similarly, combination of etomoxir with malonyl-CoA or palmitic increased viral titers by 2-fold (Fig 4F). We also quantified MDV genome copies using qPCR in MDV infected cells treated with etomoxir or clofibrate, and the results demonstrate that activation of FAO significantly reduced the viral gene copy numbers (Fig 4G).

**Fig 4:**
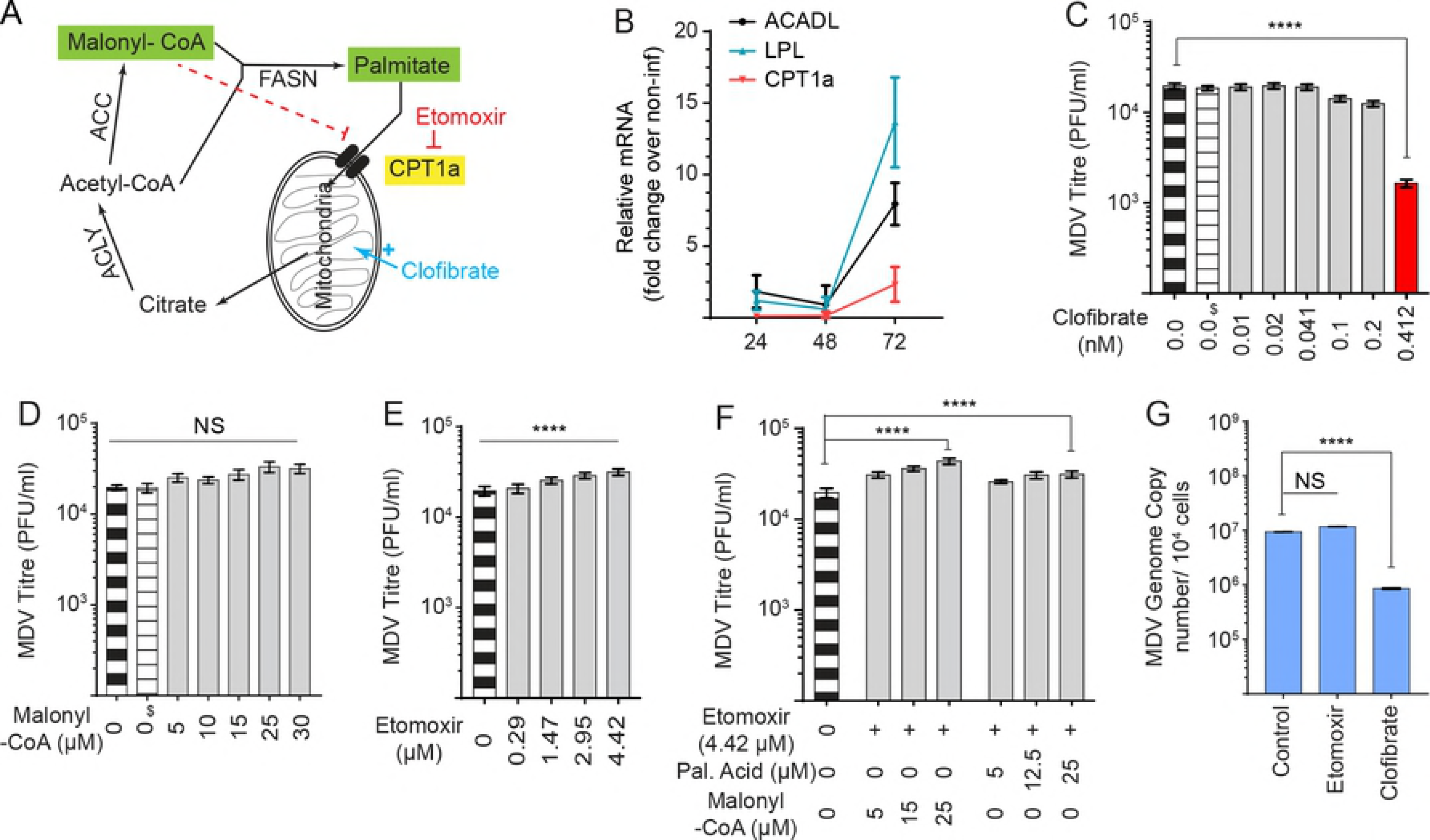
Higher virus replication occurs if the synthetized fatty acids are not used for generation of energy. Fatty acyl-CoA derivatives transported into the mitochondria undergo Fatty acid oxidation (FAO) to generate ATP, NADH and NADPH. **(A)** Schematic metabolic pathway outlines the relevant pharmacological molecules (red; etomoxir and blue; clofibrate) and the respective enzyme (yellow box) as well as metabolites (green box) studied within the FAO pathway. **(B)** Fold change gene expression comparing mock and MDV-infected CEFs at 24, 48 and 72 hpi. MDV viral titers in MDV-infected CEFs treated with **(C)** Clofibrate (0.01, 0.02, 0.04, 0.1, 0.2 and 0.41 5.9 μM), an agonist of PPAR-α, **(D)** malonyl-CoA (5, 10, 15, 25 and 30 μM) and **(E)** etomoxir (0.29, 1.47, 2.95 and 4.42 μM). **(F)** MDV-infected CEFs were treated with etomoxir (4.42 μM) with either palmitic acid (5, 12.5 and 25 μM) or malonyl-CoA (5, 15, 25 μM) and viral titer was analysed using a plaque assay. **(G)** MDV genome copy numbers per 10^4^ cells (MEQ gene with reference ovotransferrin gene) were determined using qPCR in CEFs treated with etomoxir (4.42 μM) or Clofibrate (5.9 μM). Non-parametric Wilcoxon tests (Mann-Whitney) and One-way ANOVA was used to assess normal distribution and test significance with the results shown as mean ± SD. $ indicate vehicle treated cells. **** (p < 0.0001) indicates a statistically significant difference compared to control. NS indicates no significant difference. All experiments were performed in 6 replicated and data presented is representative of all 3 independent experiments.

### MDV-induced lipogenesis results in formation of neutral lipid droplets

Lipid droplets (LD) are endoplasmic reticulum (ER) derived organelles that consists of a neutral lipids core with a phospholipid bilayer. An increase in the numbers of LD is an indication of lipogenesis. We examined if the increase in palmitic acid synthesis and excess VLCFA during MDV infection results in LD formation. Initially, Oil Red O staining was used to detect neutral lipid and hematoxylin for cell nuclei identification in mock and MDV infected CEFs at 72 hpi (Fig 5A). An accumulation of LD was observed in the MDV-infected (Fig 6Ai) compared to mock-infected cells (Fig 5Aii). To quantify the number of LD, pRB1B UL35-GFP-infected CEFs were stained with a neutral lipid dye Red LIPIDTOX (488nm), analysed by confocal microscopy and Z-stack images taken. As expected, the viral capsid protein UL35 fused to GFP was detected in the nucleus, while LD only localized within the cytoplasm (S3 Fig). A higher numbers of LD per cell were observed in MDV-infected compared to mock-infected cells (p = 0.0001) (Fig 5B). This was confirmed with an unbiased quantification of the LD using the IMARIS software (Fig 5C). Treatment of CEFs with SB 204990, or etomoxir had no effect on total LD numbers whereas treatment with TOFA and C75 decrease (p = 0.0001) the LD per cell in both mock (Fig 5D, S4 Fig) and MDV-infected cells (Fig 5E, S4 Fig). Altogether, these data indicate that MDV infection activates *de novo* lipogenesis and the synthesized fatty acids are converted into neutral lipids and stored in LD.

**Fig 5:**
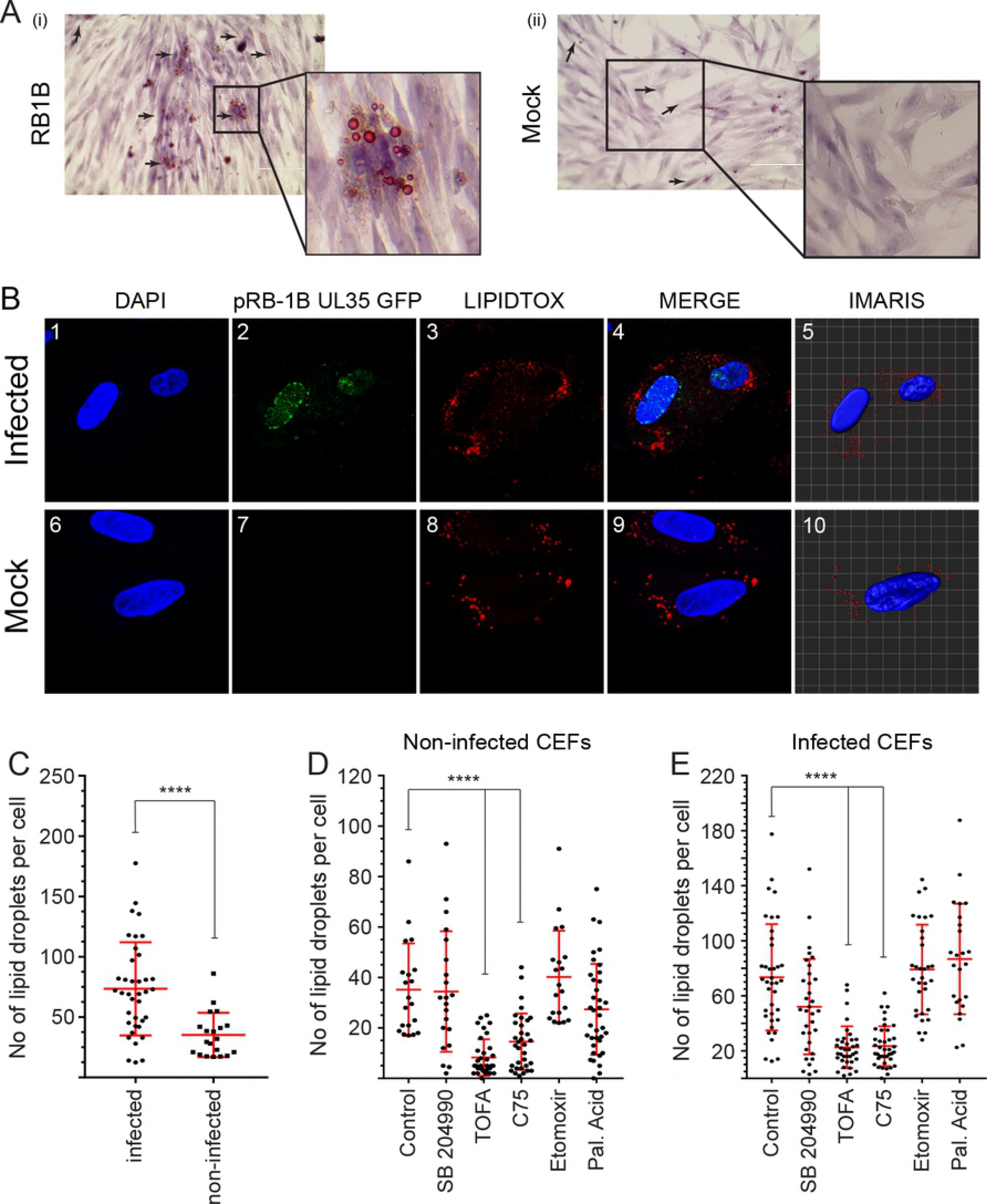
MDV-induced lipogenesis results in formation of neutral lipid droplets. **(A)** Visualisation of cytoplasmic lipids in neutral lipid droplet organelles of CEFS **(i)** infected with MDV and **(ii)** mock-infected. Cell monolayer was fixed and stained with an Oil Red O staining kit and counterstained with hematoxylin and visualized using light microscopy (magnification 10X). Black arrows indicate lipid droplets. **(B)** Confocal microscopy imaging with maximum projection of Z-stacks for each channel demonstrating nuclear and cytoplasmic distribution of pRB1B UL35-GFP virus (green) and lipid droplets (red). Mock and pRB1B UL35-GFP infected cells were fixed at 72 hpi and stained with DAPI (nuclear stain) and the neutral lipid stain LIPIDTOX-568nm. Images 5 and 10 are 3-D representative images analysed using the IMARIS software. Z-stacks were analysed using the IMARIS spot function analysis tool to quantify the relative amount of lipid droplets per cell in infected CEFs and non-infected CEFs (50-90 cells). **(C)** The numbers of lipid droplets per cell in MDV-infected and mock-infected CEFs. The numbers of lipid droplets per cell in **(D)** mock infected **(E)** MDV-infected CEFs treated with SB 204990 (3.85 μM), TOFA (1.54 μM), C75 (5.9 μM), etomoxir (4.42 μM) and palmitic acid (25 μM) are shown. **** (p < 0.0001) indicates a statistically significant difference compared to the control. All experiments were performed in duplicates and data is representative of 3 independent experiments.

**Fig 6:**
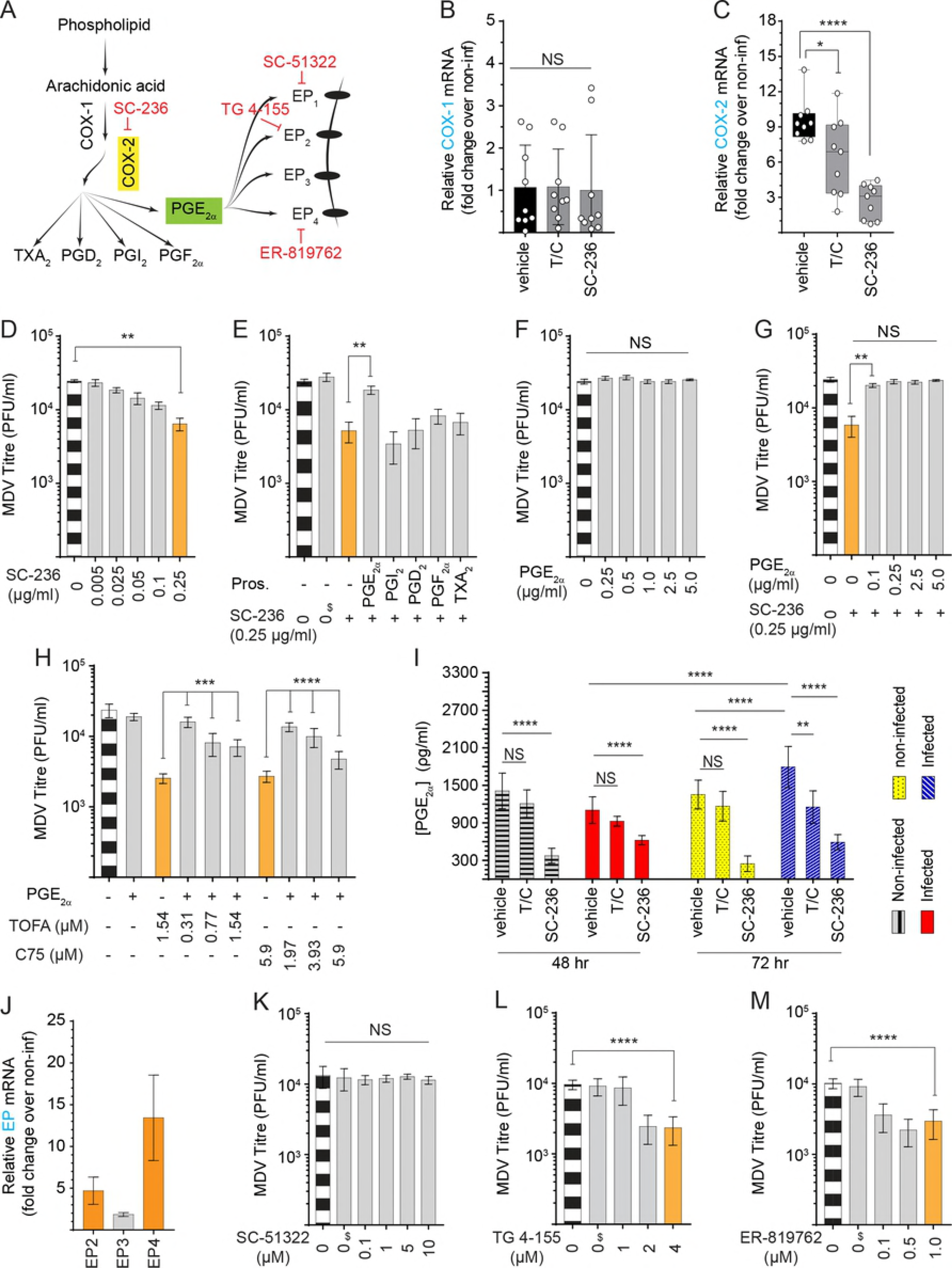
Virus-induced FA synthesis pathway activates COX-2/PGE_2α_ pathway and support virus replication through EP2 and EP4 receptors engagement. **(A)** Schematic pathway outlines the relevant pharmacological inhibitor (red) and the respective enzyme (yellow box) as well as metabolites (green box) studied within the eicosanoid biosynthesis pathway. Fold change in expressions of **(B)** COX-1 and **(C)** COX-2 in mock-infected and MDV-infected CEFs treated with T/C (TOFA; 0.5 μg/ml in combination with C75; 1.0 μg/ml), or SC-236 (0.25 μg/ml). MDV titer (PFU/ml) in the MDV-infected CEFs in the presence of **(D)** SC-236 (0.005, 0.025, 0.05, 0.1 and 0.25 μg/ml), **(E)** SC-236 (0.25 μg/ml) in combination with PGE_2α_ (5.0 μg/ml), PGI_2_ (0.5 μg/ml), PGD_2_ (0.5 μg/ml), PGF_2α_ (0.5 μg/ml), and TXA_2_ (0.5 μg/ml). MDV titer in CEFs treated with **(F)** different concentrations of PGE_2α_ (0.25, 0.5, 1.0, 2.5, and 5.0 μg/ml) **(G)** PGE_2α_ (5.0 μg/ml) in combination with SC-236 (0.25 μg/ml) and **(H)** PGE_2α_ (5.0 μg/ml) in combination with either TOFA (0.1, 0.25 and 0.5 μg/ml) or C75 (1.0, 1.2 and 1.5 μg/ml). **(I)** PGE_2_ (ρg/ml) concentrations in supernatant of mock-infected and MDV-infected CEFs were measured using an ELISA assay after 48 and 72 hpi in the presence of SC-236, T/C or no treatment. **(J)** Fold change in gene expression of PGE_2α_ receptors EP2, EP3 and EP4 in MDV-infected CEFs at 72 hpi. MDV viral titer (PFU/ml) in MDV-infected CEFs in the presence of different concentration of PGE_2α_ receptor antagonists **(K)** SC-51322 (EP1 antagonist; 0.1, 1, 5 and 10 μM), **(L)** TG 4-155 (EP2 antagonist, 1, 2 and 4 mM) and **(M)** ER-819762 (EP4 antagonist; 0.1, 0.5 and 1.0 mM). $ symbol indicates vehicle treated cells. * (p = 0.03), ** (p = 0.0022), *** (p = 0.0007) and **** (p = 0.0001) indicate statistically significant differences. NS indicates no statistical difference. Experiments were performed in 6 replicates for plaque assays and 3 replicates for Real-Time PCR and ELISA assays. The data are representative of 3 independent experiments.

### Virus-induced FA synthesis pathway activates COX-2/PGE_2α_ pathway

As shown above, AA synthesis was elevated in the MDV infected CEFs (p = 0.003) at 48 and 72 hpi (Fig 1A). AA is a substrate for the production of eicosanoids, a class of lipid mediators converted by clycooxygenase-1 (COX-1) and inducible clycooxygenase-2 (COX-2). Therefore, we investigated the importance of eicosanoid synthesis in MDV infection as shown in the schematic diagram (Fig 6A). No significant difference in COX-1 gene expression levels was observed between the mock and MDV-infected cells at 72 hpi (Fig 6B). Treatment of MDV-infected CEFs with TOFA, C75 (FA synthesis pathway inhibitors), or SC-236 (COX-2 inhibitor) did not alter the expression of COX-1 expression. However, COX-2 expression was significantly increased in the MDV-infected cells (Fig 6C) and TOFA and C75 (T/C), or SC-236 decreased (p = 0.0007) COX-2 mRNA expression, suggesting that FA synthesis pathway is involved in activation of COX-2 expression. Treatment of MDV-infected cells with non-toxic concentrations of a COX-2 inhibitor (SC-236) diminished MDV replication in a dose dependent manner. Specifically, 4-fold reduction in MDV titers was observed in CEFs treated with SC-236 (p = 0.0022) (Fig 6D). We demonstrated that added PGE_2α_ is the only prostaglandin that can rescue the inhibitory effects of SC-236 on MDV titer (Fig 6E), while exogenous PGE_2α_ did not alter MDV titer (Fig 6F). A concentration as low as 0.1 μg/ml of PGE2α sufficed to rescue the inhibitory effects of SC-236 on MDV titer (Fig 6G). Strikingly, exogenous PGE_2α_ also restored the inhibitory effects of TOFA and C75 on virus titer (Fig 6H), suggesting that the inhibitory effects of FA synthesis pathway inhibitors is dependent on inhibition of PGE_2α_ synthesis. When CEFs were treated with low concentrations of TOFA or C75, exogenous PGE_2α_ fully restored MDV replication (p = 0.0007). However, PGE_2α_ only partially recovered MDV titer when high concentrations of TOFA or C75 were used (Fig 6H). To determine if PGE_2α_ synthesis is increased in MDV infected cells, we measured PGE_2α_ in the supernatants of mock and MDV infected cells at 48 and 72hpi by ELISA (Fig 6I).

Higher concentrations of PGE_2α_ were detected in supernatants of MDV-infected cells compared to mock-infected cells at 72hpi (p = 0.0001). Interestingly, inhibition of FA synthesis pathway by TOFA and C75 reduced PGE_2α_ release from MDV-infected cells (p = 0.0022), but not the mock infected cells at 72hpi. The results indicate that the treatment with TOFA and C75 only reduces MDV-induced PGE_2α_ synthesis, while physiological levels of PGE_2α_ are still produced in treated cells. While, SC-236 significantly inhibited PGE_2α_ production from both the MDV-infected (p = 0.0001) and mock-infected (p = 0.0001) cells at 48 and 72hpi (Fig 6I). Taken together, our results suggest that virus-induced FA synthesis pathway is critical for the increased PGE_2α_ synthesis in the MDV-infected cells.

### Engagement of EP2 and EP4 receptors are involved in MDV replication

Four different types of PGE_2α_ receptors have been identified in humans are EP1, EP2, EP3, and EP4. Chicken EP2, EP3, and EP4 receptors have been cloned and characterized [31, 32] while chicken EP1 receptor has not yet been identified. Analysis of gene expressions in the MDV-infected and mock-infected CEFs demonstrated that the EP2 and EP4 receptors were significantly up-regulated in cells infected with MDV (Fig 6J), suggesting that PGE_2α_ may support MDV replication through an EP2 or/and EP4 mediated mechanisms. We used receptor antagonists for EP1 (SC-51322), EP2 (TG 4-155) and EP4 (ER-819762). As expected, the EP1 receptor antagonist (SC-51322; 0.1, 1, 5 and 10 μM) had no effect on MDV replication. In contrast, TG 4-155 (2 and 4 μM) and ER-819762 (0.1, 0.5 and 1.0 μM) inhibited MDV infection (p = 0.0001), confirming that PGE_2α_ synthesis supports MDV replication through engagement of the EP2 and EP4 receptors. Taken together, our data indicate that virus-induced FA synthesis enhances PGE_2α_ synthesis and that this process is dependent on the EP2 and EP4 receptors in the MDV-infected cells.

## Discussion

Viruses have evolved strategies to target and modulate lipid signalling, synthesis and metabolism in the host cells and provide an optimal micro-environment for viral entry, replication and morphogenesis. Interestingly, even two related viruses such as HSV-1 and HCMV drive host cells to achieve distinct metabolic programs. While HCMV increased *de novo* lipid synthesis, HSV-1 enhanced the synthesis of pyrimidine nucleotides [7]. Other viruses can also modulate common lipid metabolism including upregulation of FA synthesis pathway, providing building blocks for different lipids. Replication of different viruses may require activation of distinct lipid pathways and source of energy. MDV is an *alphaherpesvirus* that infects chickens and causes a deadly lymphoproliferative disease. In addition to transformation of CD4+ T cells, MDV causes atherosclerosis by disturbing the lipid metabolism in the infected chickens [33], which can be inhibited by vaccine-induced immunity [29, 34]. Surprisingly, little is known about the processes involved in disturbance of lipid metabolism in MDV infection. Our lipidomic analysis of MDV-infected CEFs demonstrates that MDV infection enhances FA synthesis and promote synthesis of AA, the prostaglandin precursor. Here, we demonstrate that MDV hijacks host metabolic pathways to provide essential macromolecular synthesis to support infection and replication. *De novo* FA synthesis generates the metabolic intermediate acetyl-CoA, malonyl-CoA and finally palmitate. Targeted inhibition studies against the enzymes involved in FA synthesis during infection have yielded an alternative understanding of alteration of lipid metabolism in infections with HCMV [35], EBV [22], HCV [36, 37]. The first step towards FA synthesis is the conversion of citric acid into acetyl-CoA by direct phosphorylation of ACLY. Hepatitis B virus (HBV) and HCMV infections activate the expression of ACLY [38, 39], however the enzymatic activity of ACLY is not critical for virus-induced lipogenesis. Virus-infected cells can use glucose carbon for FA synthesis by ACLY from citrate generated in mitochondria and acetyl-CoA synthetase short-chain family member 2 (ACSS2) from acetate [40]. In our study, blocking ACLY activity via small pharmacological inhibitors had no effect on MDV replication and genome copies. This finding suggests that acetate can be produced in avian cells via other endogenous mechanism under rich medium conditions, which would mask the effects on viral replication. The subsequent step involves the conversion of acetyl-CoA into malonyl-CoA by acetyl-coA carboxylase (ACC) and finally elongation by utilizing both acetyl-CoA and malonyl-CoA coupled to the multifunctional fatty acid synthase (FASN) to make palmitic acid. Our results demonstrate that blocking ACC and FASN activity significantly reduces MDV replication, suggesting that MDV preferentially modulates FA synthesis pathway to generate a variety of lipids which contributes to several key cellular processes. Our data confirm that the inhibitory effects of FA synthesis pathway inhibitors on MDV replication could be overcome by the addition of palmitic acid, a metabolite downstream of FASN in the FA synthesis pathway. This indicates that inhibition of MDV by FA synthesis inhibitors was not simply detrimental to the cell but was essential for the production of infectious virus. Future studies are required to examine the mechanism involved in activation of FA synthesis by MDV which can lead to induction and accumulation of fatty acids.

Palmitic acid contributes essential carbons in synthesis of VLCFA that can be subsequently utilized in synthesis of various lipids required for membrane biosynthesis and lipid droplet formation. Our results demonstrate that VLCFA are increased in MDV infection. Alternatively, β-oxidation of fatty acids within the mitochondria can generate energy and macromolecular precursors to support cellular activity and fuel viral replication. It has been shown that the synthesis and mitochondrial import of fatty acids, in which β-oxidation generate ATP production, are essential for replication of vaccinia virus [11]. This can be explained by the fact that vaccinia virus does not require glycolysis for replication and thus depends on the energy generation via β-oxidation. Our results demonstrate that β-oxidation is activated in the MDV-infected cells and palmitic acid is converted into energy in the mitochondria. However, we could demonstrate that limiting energy derived from mitochondrial β-oxidation during lytic viral infection had no detrimental effect on MDV replication. In fact, inhibition of β-oxidation rather increased virus titers, suggesting that MDV-infected cells obtain their energy from other sources. This could be explained by the facts that some herpesviruses encode metabolic enzymes to support essential functions independent of mitochondria derived energy carriers [1–4]. Alternatively, MDV could induces glycolysis and thus does not require energy generated via β-oxidation as observed for other herpes viruses [7, 35]. Consequently, activation of lipolysis (mitochondrial β-oxidation) decreased virus titers and genome copies, confirming that β-oxidation is not beneficial for MDV replication. This indicates that the generated fatty acids are utilized for activation of other lipid pathways, which are crucial for viral replication.

It is known that elongation of palmitic acid in lipogenesis contributes to a total cellular pool of VLCFA which are essential components for the initiation of key cellular processes such as membrane lipid synthesis, generation of lipid droplets and eicosanoid synthesis [41]. It has been shown that the assembly of some viruses such as HCMV are highly dependent on induction of VLCFA from FA synthesis [23]. Similarly, VLFCA contribute to IAV envelope formation in a strain dependent manner [42]. Lipid droplets are classically defined as organelles with stored neutral lipids and some reports suggest that lipid droplets are a site for replication and assembly of some viruses. However the functional relationship of lipid droplets to cellular processes are not well defined [43]. Infection with a range of pathogens have been demonstrated to induce lipid droplet formation including bacteria, specifically *M. tuberculosis* [44] and *M leprae* [45], as well as protozoan infection such as *Trypanosoma cruzi* [46] in Chagas disease. Viral infection including HCV and its relationship to lipid droplet formation has been extensively documented [47]. It has been demonstrated that the HCV core protein interacts with lipid droplet for virion assembly [48]. Similarly, infection with HCV [48], DENV [49] and rotaviruses [50] promoted the formation of lipid droplets. Similarly, our data demonstrate that MDV infection increases lipid droplet formation; however, the exact role of lipid droplets in MDV infection is unknown. Formation of lipid droplets in MDV infection was dependent on FA synthesis pathway as FA synthesis pathway inhibitors reduced the numbers of lipid droplets. Lipid droplets are a significant source of triglyceride-derived arachidonic acid (AA) which can be converted to eicosanoids such as prostaglandins. Upon stimulation, AA is released from lipid droplets and metabolised into PGE_2α_ by the cyclooxygenase enzymes COX-1 and COX-2. In this study, we observed that MDV promotes lipid droplet formation, and synthesis of phospholipid and AA. The upregulation of COX-2 but not COX-1 transcripts in MDV-infected cells confirms that COX-2/PGE_2α_ pathway are induced by the virus. Some viruses such as EBV [51] promote viral replication and dissemination by supressing PGE_2α_ biosynthesis. In contrast, HCMV [17, 52], KSHV [20] and MVA [16] activate PGE_2α_ release in response to infection. Here, we demonstrate that MDV upregulate COX-2 expression and PGE_2α_ synthesis, which is highly dependent on the FA synthesis pathway. To our knowledge, this is the first report demonstrating that a viral infection increases COX-2/PGE_2α_ pathway via induction of FA synthesis pathway, and demonstrate that the inhibitory effects of FA synthesis pathway inhibitors on virus replication could be recovered by exogenous PGE_2α_. The proposed model for the role of FA synthesis pathway and COX-2/PGE2_2α_ pathways in MDV infection is summarized in Figure 7. Taken together, our results demonstrate that virus-induced FA synthesis pathway enhances PGE_2α_ synthesis which support MDV infectivity through EP2 and EP4 receptors.

**Fig 7:**
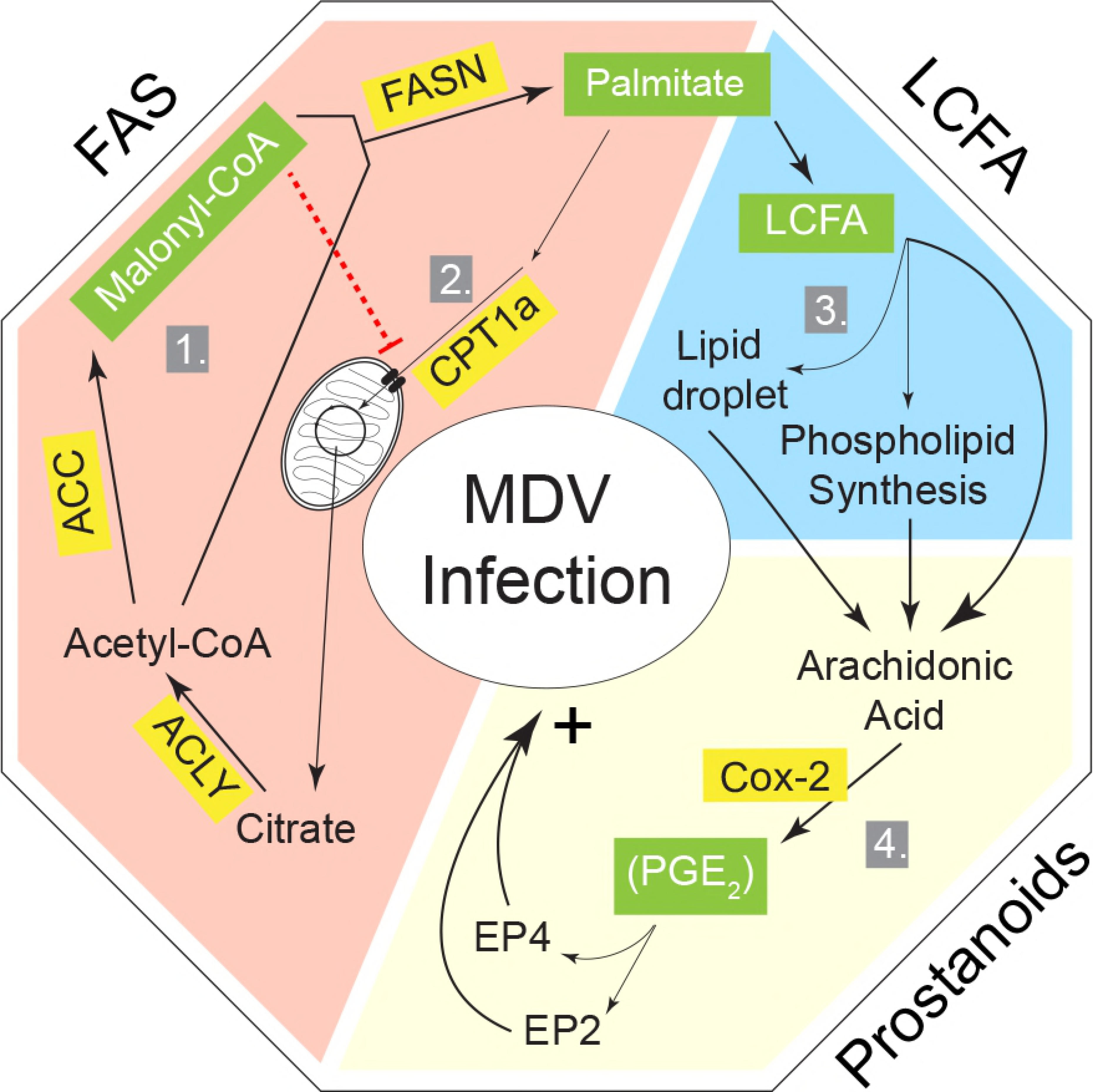
Schematic representation of the proposed model by which MDV activates FA synthesis pathway and COX-2/PGE2 pathways and promote virus replication through EP2 and EP4 receptors engagements. MDV systematically modulates cellular lipid metabolic pathways to support its replication through FA synthesis pathway and COX-2/PGE2 pathways. The three major phases of lipogenesis are grouped under FA synthesis pathway, VLCFA and Prostanoids. **1.** Infection of CEFs with MDV results in an increase of *de novo* FA synthesis pathway, which is required for MDV replication. **2.** Palmitic acid biosynthesized in FA synthesis pathway can be utilized in the mitochondria by FAO (β-oxidation) to regenerate energy intermediates, however this pathway is not required for virus replication. **3.** Elongation of palmitic acid results in biosynthesis of VLCFA which can be stored in lipid droplets organelles, utilized for phospholipid or eicosanoid synthesis **4.** Subsequently, PGE_2α_ are synthesized enzymatically from arachidonic acid. Signalling of PGE_2α_ is mediated through its EP2 and EP4 receptors in MDV-infected cells. Specific enzymes are highlighted in yellow and major lipid metabolites are highlighted in green.

## Materials and Methods

### Ethics Statement

Ten day old mixed sex SPF embryonated chicken eggs which were purchased from Valo (Valo Biomedia GmbH), and were used to generate primary chicken embryonic fibroblast cells (CEFs). All embryonated chicken eggs were handled in strict accordance with the guidance and regulations of European and United Kingdom Home Office regulations under project licence number 30/3169. As part of this process the work has undergone scrutiny and approval by the ethics committee at The Pirbright Institute.

### CEFs culture and virus preparations

CEFs were generated from mixed sex SPF Valo eggs (Valo Biomedia GmbH) incubated in a Brinsea Ova-Easy 190 incubator at 37°C until 10 days *in ovo.* CEFs were seeded at a rate of 1.5 × 10^5^ cells /ml in 24 well plates with growth medium (E199 supplemented with 10% TBP, 5% FCS, 2.8% SQ water, amphotericin B (0.01%), Penicillin (10 U/ml) and Streptomycin (10 μg/ml)) and incubated overnight (38.5°C at 5% CO_2_). Next day, 80% confluent monolayer was observed and growth medium was removed and replaced with maintenance medium (E199 supplemented with 10% TBP, 2.5% FCS, 3.5 % SQ water, amphotericin B (0.01%), Penicillin (10 U/ml) and Streptomycin (10 μg/ml). RB1B (MDV; serotype 1) and RB-1B UL35-GFP virus expressing GFP fused to the UL35 capsid protein were prepared in CEFs.

### Reagents and antibodies

Chemicals: SB 204990 (Thermo Fisher Scientific, Paisley, UK), TOFA, C75, Clofibrate, Palmitic acid, SC-236 (Sigma-Aldrich, Dorset, UK), PGD_2_, PGI_2_, PGE_2α_ (Cambridge Bioscience, Cambridge, UK), SC-51322, TG-4-155 and ER-819762 (Bio-Techne Ltd., Abingdon, UK) were all reconstituted in DMSO. TXA_2_ (Cambridge Bioscience, Cambridge, UK) was reconstitute in ethanol. PGF_2α_ (Cambridge Bioscience, Cambridge, UK), etomoxir and malonyl-CoA (Sigma-Aldrich, Dorset, UK) were reconstituted in E199 medium.

### Cells and MDV Infection

**Metabolomics:** CEFs were either mock infected or with the very virulent RB1B strain (100 pfu per 1.5 × 10^5^ cells) in triplicates and harvested at 48 and 72 h post infection (hpi). The cells were washed, counted and after protein quantification using Bradford assay, the samples were sent for Metabolom analysis using GCxGC-MS (Target Discovery Institute, University of Oxford). In brief, the cells were homogenized using bead beater in methanol/water (1:1), and then t-butyl methyl ether was added for phase separation. The organic phase was dried under vacuum, while methanol was added to the remaining sample and mixed in bead beater. After incubation at −80°C for 1 hr, the phase separation occurred after centrifugation, and liquid layer was collected and dried under vacuum. Methoxyamine and MSTFA (1% TMSCI) were added to the dried samples and subsequently injected for analysis by GC/GC-MS. The method used in this experiment is designed to detect 155 different metabolites including lipids and amino acids. The lipid profile of mock and MDV-infected cells were analysed in biological triplicates with up to six technical replicates per biological replicate. The data were adjusted and normalized based on protein content. Virus infection did not change the size of the cells as determined by microscopy.

**Viral plaque analysis:** Cells were fixed (1:1 acetone:methanol) and blocked with blocking buffer (PBS + 5% FCS) for 1 hr at RT, subsequently incubated with anti-gB mAb (HB-3) and then with horse radish peroxidase-conjugated rabbit anti-mouse Ig. After development of the plaques using AEC substrate, the cells were washed with super Q water and viral plaques were counted using light microscopy.

**Determining non-toxic concentration of the inhibitors:** To identify non-toxic concentrations of the chemicals, mock-infected and MDV-infected CEFs were exposed to the chemicals or vehicles and cell morphology and adherence/confluency were monitored under light microscopy at different time points post treatment. Moreover, CEFs were trypsinized, stained with 7-AAD (BD Bioscience, Oxford, UK) and acquired using a MACS quant flow cytometry and FloJo software for analysis of the data (S1 Fig). Non-toxic concentrations of the inhibitors and chemicals were selected based on flow cytometry data and confluency.

### qPCR to amplify MDV genes

DNA samples were isolated from 5 × 10^6^ cells using the DNeasy-96 kit (Qiagen, Manchester, UK), according to the manufacturer’s instructions. A master-mix was prepared: primers Meq-FP and Meq-RP (0.4 μM), Meq probes (0.2 μM), *ovo* forward and reverse primers (0.4 μM), and *ovo* probe (0.2 μM, 5′Yakima Yellow-3′TAMRA, Eurogentec) and ABsolute Blue^®^ q-PCR Low Rox master-mix (Thermo Fisher Scientific, Paisley, UK). A standard curve generated for both Meq (10-fold serial dilutions prepared from plasmid construct with Meq target) and *ovo* gene (10-fold serial dilutions prepared from plasmid construct with *ovo* target) were used to normalise DNA samples and to quantify MDV genomes per 10^4^ cells. All reactions were performed in triplicates to detect both *Meq* and the chicken ovotransferrin (ovo) gene on an ABI7500^®^ system (Applied Biosystems) using standard conditions. MDV genomes were normalised and reported as viral genome per 10^4^ cells.

### Real-Time Polymerase Chain Reaction (RT-PCR)

*RNA extraction and cDNA:* Total RNA was extracted from mock and MDV infected CEFs using TRIzol (Thermo Fisher Scientific, Paisley, UK) according to the manufacturer’s protocol and treated with DNA Free DNase. Subsequently, 1 μg of purified RNA was reverse transcribed to cDNA using Superscript^®^ III First Strand Synthesis kit (Thermo Fisher Scientific, Paisley, UK) and oligo-dT primers according to the manufacturer’s recommended protocol. The resulting cDNA was diluted 1:10 in DEPC treated water.

*SYBR green RT-PCR:* Quantitative real-time PCR using SYBR Green was performed on diluted cDNA using the LightCycler^®^ 480 II (Roche Diagnostics GmbH, Mannheim, GER) as previously described. Briefly, each reaction involved a pre-incubation at 95 °C for 5 min, followed by 40 cycles of 95°C for 20 sec, 55°C-64°C (T_A_ as per primer) for 15 s, and elongation at 72°C for 10 s. Subsequent melt curve analysis was performed by heating to 95°C for 10 sec, cooling to 65°C for 1 min, and heating to 97°C. Primers sequences and accession numbers are outlined in S1 Table. Relative expression levels of all genes were calculated relative to the housekeeping gene β-actin using the LightCycler^®^ 480 Software (Roche Diagnostics GmbH, Mannheim, GER). Data represent mean of 6 biological replicates.

### Prostaglandin E_2α_ ELISA

PGE_2α_ was quantified using a colorimetric assay (R&D Systems, Abingdon, UK) based on competition between unlabelled PGE_2α_ in the sample and a fixed amount of conjugated PGE_2α_. The assay was performed according to assay kit manufactures recommendation.

In brief, CEFs were either mock or infected with RB1B in the presence/absence of TOFA (1.54 μM) and C75 (5.9 μM) or SC-236 (0.25 μg/ml) and incubated (38.5°C, 5% CO_2_) for 48 or 72 hpi. The supernatants were collected and diluted (1:100) in Calibrator Diluent RD5-56. PGE_2α_ detection antibody was added to the wells which were pre-coated with primary anti-PGE_2α_ antibody and incubated for 1 hr at RT. Next, PGE_2α_ conjugates was added and incubated for an additional 2 hr at RT. The wells were subsequently washed and developed by incubating substrate buffer for 30 min at RT. The assay was measured at OD_450_ nm and concentrations of PGE_2α_ were determined against a standard curve.

### Determination of Oxygen consumption rate (OCR) and Extracellular acidification rate (ECAR)

CEFs were seeded at a rate of 2.0 × 10^4^ cells per well in an XFp 8-well V-3 PET tissue culture mini-plate (Seahorse Bioscience, UK) in triplicates and incubated overnight at 38.5°C. Next day, cells at 80% confluency were pre-treated with etomoxir (4.42 μM) or vehicle control and then either mock or infected with RB1B (100 pfu per 1.5 × 10^5^ cells) for an additional 24 h. Oxygen consumption rates (OCR) was measured every 6 min using the Seahorse Bioscience XFp analyzer (Seahorse Bioscience, UK) from 16-30 hpi at 38.5°C.

### Oil Red O staining

Lipid droplets were stained in CEFs mock or infected with RB1B in 6 well-plates. In brief at 72 hpi, cell monolayer was washed with PBS and fixed with 4% formaldehyde for 30 min at RT. Plates were subsequently stained with Oil Red O solution for 30 min at RT followed by a wash in PBS. Plates were counterstained with hematoxylin for 3 min at RT followed by a wash with super Q water. Plates were visualized and imaged using a light microscope and the pictures were processed using Adobe Photoshop software.

### Fluorescence confocal microscopy

CEFs were seeded in 24 well plates that contained 12 mm diameter round coverslips at a rate of 1.0 × 10^5^ cells per well. At 72h post mock or infection with the pRB1B UL35-GFP virus in the presence/absence of SB 204990, TOFA, C75 or etomoxir, the samples were prepared for imaging. In brief, mock or infected CEFs were fixed with 4% formaldehyde for 30 min at RT and washed twice with PBS. Cells were subsequently incubated with HCS LipidTox Red Neutral lipid stain (1:1000 in PBS; 568 nm) at RT for an additional 45 min. Cells were washed twice with PBS and nuclei were labelled with DAPI. Coverslips were mounted in vectashield mounting medium for fluorescence imaging. Cells were viewed using a Leica SP2 laser-scanning confocal microscope and optical sections recorded using either the 663 or 640 oil-immersion objective with a numerical aperture of 1.4 and 1.25, respectively. All data were collected sequentially to minimize cross-talk between fluorescent signals. The data are presented as maximum projections of z-stacks (23-25 sections; spacing 0.3 mm). Maximum projections of z-stacks were analysed using IMARIS (Bitplane Scientific Software). Ninety infected cells and 40 mock-infected cells were analysed and their LipidTox-labelled Neutral lipid containing organelles were detected with the spot function of IMARIS. Images were processed using Adobe Photoshop software.

### Statistical Analysis

All data are presented as mean ± standard deviation (SD) from at least three independent experiments. Quantification was performed using Graph Pad Prism 7 for windows. The differences between groups, in each experiment, were analysed by non-parametric Wilcoxon tests (Mann-Whitney) or by Kruskal-Wallis test (One-way ANOVA, non-parametric). Results were considered statistically significant at *P* < 0.05 (*).

## Acknowledgments

Analysis of metabolomics data was initially performed by Metabolon (London, UK). Our special thanks to Dr. Venugopal Nair (The Pirbright Institute, UK) and Dr. Miriam Pedrera (The Pirbright Institute, UK) for providing us with the pRB1B UL35-GFP virus and excellent support in generating viral stocks.

---

**S1 Fig:**Analysis by MACS quant demonstrating percentages of 7AAD negative CEFs, representing live cells, after 72 h treatment with small pharmacological inhibitors including **(A)** SB 204990, **(B)** TOFA, **(C)** C75, **(D)** Palmitic Acid, **(E)** Malonyl-CoA, **(F)** Etomoxir, **(G)** Clofibrate, **(H)** SC-236, **(I)** ER-819762, **(J)** SC-51322 and **(K)** TG-4-155. Bar graphs with single bars in black represent concentrations of the inhibitors which did not induce cell death.

**S2 Fig:** OCR (pmoles/min) of mock-infected CEFs treated with either etomoxir (4.42 μM) or vehicle are shown for 12 h. All experiments were performed in triplicates and data is representative of 3 independent experiments.

**S3 Fig:** Analysis by IMARIS, using the co-localization function, to visualize neutral lipid droplet organelles (red) formation and relative position to the pRB1B UL35-GFP virus capsid protein (Green). **(A)** IMARIS software spot function visualisation of lipid droplet organelles (Red) to the small capsid protein (UL_35_) with the GFP tag (Green). **(B)** Magnification of the first picture where co-localization analysis was performed

**S4 Fig:** Visualisation of cytoplasmic lipids in neutral lipid droplet organelles by confocal microscopy imaging with maximum projection of Z-stacks for each channel demonstrating nuclear and cytoplasmic distribution of pRB1B UL35-GFP virus (Green) and lipid droplets (Red) in CEFs treated with the small pharmacological inhibitors **(A)** SB 204990, **(B)** TOFA, **(C)** C75, **(D)** etomoxir and **(E)** Palmitic Acid.

**S1 Table:** List of primers used in this study.

